# The petrous bone contains high concentrations of osteocytes: one possible reason why ancient DNA is better preserved in this bone

**DOI:** 10.1101/2022.05.20.492830

**Authors:** Jamal Ibrahim, Vlad Brumfeld, Yoseph Addadi, Sarah Rubin, Steve Weiner, Elisabetta Boaretto

## Abstract

The characterization of ancient DNA in fossil bones is providing invaluable information on the genetics of past human and other animal populations. These studies have been aided enormously by the discovery that ancient DNA is relatively well preserved in the petrous bone compared to most other bones. The reasons for this better preservation are however not well understood. Here we examine the hypothesis that one reason for better DNA preservation in the petrous bone is that fresh petrous bone contains more DNA than other bones. We therefore determined the concentrations of osteocyte cells occluded inside lacunae within the petrous bone and compared these concentrations to other bones from the domestic pig using high resolution microCT. We show that the concentrations of osteocyte lacunae in the inner layer of the pig petrous bone adjacent to the otic chamber are about three times higher than in the temporal bone, as well as the cortical bone of the femur. The sizes and shapes of the lacuna in the inner layer of the petrous bone are similar to those in the femur. We also confirm that the petrous bone lacunae do contain osteocytes using a histological stain for DNA. We therefore conclude that one possible reason for better preservation of ancient DNA in the petrous bone is that this bone initially contains up to three times more DNA than other bones, and hence during diagenesis more DNA is likely to be preserved. We also note that the osteocytes in the inner layer of the petrous bone may have a function in hearing.

## Introduction

The possibility of extracting genetic information from fossil material was first demonstrated by Higuchi (1). Many subsequent studies have confirmed the presence of ancient DNA in fossil bones (2-5), but have also shown that contamination and preservation are severe problems (6, 7). In 2014 Gamba *et al*. made a remarkable discovery: aDNA is better preserved in the petrous bone that surrounds the otic chamber as compared to other bones (8). This has been confirmed by other studies on fossil petrous bones (9-11). The results obtained from analyzing aDNA in fossil petrous bones has and is contributing significantly to our understanding of ancient genomes, and in particular those of hominins. For example, successful retrieval of aDNA in the petrous bone was carried out for the European Late Upper Paleolithic around 13.3k years ago from western Georgia and 13.7k years ago from Switzerland, and from a Mesolithic male from western Georgia 9.7k years ago (12). A study of 230 ancient Eurasians who lived between 6500-300BC was made possible by obtaining excellent aDNA amounts from petrous bones (13). Genome-wide data extracted from 16 prehistoric petrous bones of Africans hominins unraveled episodes of natural selection in southern African populations (14). However, the reasons why DNA is better preserved in the petrous bone compared to other bones are not known. It has often been assumed that this better preservation is somehow related to the unusually high density of the petrous bone (8). Another hypothesis is that bones that are most likely to preserve DNA are those with the highest amounts of endogenous DNA in vivo, namely bones with the highest concentrations of osteocytes (15).

The human petrous bone embryology, structure and extreme hardness were studied for more than a century (16). Meyer described the anatomy of the otic capsule including the layered petrous bone (16). Doden and Halves documented mineralization and structural differences in the layers of the petrous bone based on microradiography and circularly polarized light (17). They provided information for distinguishing the three layers: the layer closest to the otic chamber is referred to as the endosteal/endochondral layer; the next layer is the periosteal layer which is divided into inner (closer to the otic chamber) and outer (furthest from the otic chamber) layers (17). Doden and Halves showed that the inner two layers are more mineralized than the outer layer and contain localized discontinuous regions that are hypermineralized. In a study of quantum bone remodeling in young pigs, Sorensen *et al* showed by microradiography that remodeling in the otic capsule and its surrounding bone is absent (18).

Mendoza and Rius analyzed osteocyte lacunae from 50 otic capsule specimens from humans between 18 and 80 years old. They documented osteocyte lacunae vitality after Hematoxylin and Eosin staining (H&E) of bone sections of the endochondral and periosteal layers of the petrous bone. Their results show high rates of osteocyte necrosis in the endochondral layer (19). However in the periosteal layer the nuclei of osteocytes were always present (19). Here we address the Geigl and Grange hypothesis that one possible reason that DNA is relatively well preserved in fossil petrous bone is because the fresh petrous bone contains significantly more DNA than other bones (15). Andranowski measured DNA yield from different bone tissues and found a correlation between the number of osteocyte lacuna number and DNA yields (20). We do this by quantifying the concentration of bone cell cavities, namely osteocytic lacunae (OL) in the periosteal layer, and we show that the lacunae contain nucleated osteocytes. We show that in pigs there are up to 3 times more osteocytes in the inner layer of the petrous bone compared to the temporal bone and the femur. These petrous bone osteocytes are therefore likely to be a concentrated source of DNA embedded in bone and hence this could be one explanation for improved preservation of aDNA in the petrous bone.

## Materials and Methods Materials

### Osteocyte lacuna counting

Freshly sacrificed adult pigs (*Sus scrofa*) 7 months old (L2) and 5 months old (L3) were supplied by Lahav Clinical Research Organization (C.R.O), The Institute of Animal Research, Israel. All C.R.O. research studies are coordinated and approved by the national ethics committee for animal studies in Israel. All facilities and activities of Lahav C.R.O. are accredited in accordance with GLP and ISO9001 (2015). Samples extracted from the skulls included petrous bones which were analyzed from two regions (layers) based on the degree of calcification: outer periosteal layer (OP2 from skull L2 and OP3 from skull L3), and inner periosteal layer (IP2 and IP3). From the same skulls two temporal bone samples were obtained close to the cochlear regions (T2 and T3). Two cortical bone samples from the proximal cortex of the femurs of two different adult pigs (F1 and F2; 7 months old supplied by Lahav C.R.O) were stored in a freezer for 6 years. All cleaned samples were stored in a -20 °C freezer prior to analysis.

### Nuclei staining

A freshly sacrificed adult pig (5 months old) skull obtained from Lahav C.R.O. (Israel) was used to extract the left petrous and temporal bones. In addition, six petrous bones were extracted from 3 freshly sacrificed wild-type (ICR CD-1) female mice. All mice experiments were performed according to the Animal Protection Guidelines of Weizmann Institute of Science, Rehovot, Israel. The mice experiments were pre-approved by the relevant Weizmann Institute Institutional Animal Care and Use Committee IACUC (#01390120-1, 01330120-2, 33520117-2).

All reagents for bone clearing were purchased from Sigma Aldrich®, and DRAQ5 for nuclear DNA staining was purchased from eBioscience™ (cat. No. 65-0880-92).

## Methods

### Characterizing lacunae concentration and morphology

#### Sample preparation

The two pig skulls (L2 and L3) were dissected and cleaned immediately after sacrifice. Cutting and isolation of the skull bone using a manual saw started 3 days after storing at 4°C. For each skull, a sagittal cut was first made at the center of the lateral end of the skull, halving the occipital protuberance and separating the occipital condyles.

The cut continued until it reached the parietal bone dorsally and the palatine bone ventrally. Then second and third sagittal cuts were made on each side of the skulls between the zygomatic arch and the external acoustic opening at an angle of 50° towards the parietal bone dorsally and the palatine bone ventrally. Each part of the skull that was extracted, included one petrous bone, part of the parietal bone, foramen magnum, the foramen orbitorotundum, the bony tentorium cerebelli, the cranial cavity, one side of the paracondylar process and one side of the tympanic bulla (S.I. Movie 1). After isolation of the bones, all soft tissues were discarded, and the bones were washed with double distilled (DD) water. Using a surgical blade, the meninges was cut with the help of forceps to enable release of the petrous bone. Then the skull piece was mounted on a vice and the bone was forced wide open in the sagittal direction to release the petrous bone, without causing any deformation or breakage of the petrous bones. The cortical bones of two pig femurs were sampled from the cortex of proximal extremities.

All bone samples were washed three times in PBS solution with gentle shaking for 30 min each time. Then the bones were fixed overnight with 3% PFA, and the next day the bones were lyophilized overnight. After dehydration, the bone samples were then embedded in Epon (VersoCit-2kit, Struers, USA) and allowed to polymerize at room temperature for about 15 minutes. The embedded samples were then cut precisely using a water-cooled diamond disk (Isomet, Buehler Ltd. USA). The cut was made at the preferred location and in a specific orientation to include all layers of the pig petrous bone following Sorenson characterization of the petrous bone layers (18). The petrous bone samples were cut in the transverse plane with slight tilting to include the vestibular region and some cross-sections of the semicircular canals. The inferior sections of the petrous bones were used in this study. The cut section surfaces were then ground manually starting with grid 350 paper for 3 minutes and increasing gradually to grid 600 paper for 3 minutes each. The sections were then polished using a 1μm diamond suspension for 5 minutes. All polished sections were imaged using a light microscope in reflected light (Nikon Eclipse, USA).

#### SEM imaging

The polished surfaces were then imaged using a back scattered electron detector in a Phenom SEM (Phenom World, USA). Tiles of images were obtained automatically at 1500x magnification and then automatically stitched together in automated image map (AIM).

#### Micro CT imaging

At first, low resolution (voxel size 12 μm) CT scans were obtained and analyzed to enable selection of regions of interest for the high resolution scans. Then high resolution micro-CT scans were obtained with a small voxel size to detect and count osteocyte lacunae within small volumes of bone. The high resolution μ-CT scans were performed under the same scanning conditions: objective 0.4x, voltage=60kV, power=5W. The automatically reconstructed volumes had an average bone volume of 0.4±0.1mm^−3^, and voxel size of 0.85±0.1μm (n=8). This should be sufficiently small to accurately estimate OL volumes. All high resolution μ-CT scans were imaged over 360 degrees using 1601 projections. We do note however that the two samples of the temporal bone were embedded in large volumes of plastic. This resulted in a reduction in the signal intensity, and hence an increase in the noise of the data.

#### Lacunae counting

We used two methods for image analysis. Firstly, we used the FIJI (NIH, USA) software for traditional segmentation by applying the mean thresholding function (21). The same threshold was used for all samples. Secondly, we segmented the images by applying a deep learning model for image segmentation (DLM). For this we used the Dragonfly image analysis and deep-learning software (version 2022.1 build 1249, Objects Research Systems, Montreal, QC, Canada). For the DLM we used the Snap Tool for manually segmenting the OL and canals. Everything else is considered background. To train the DLM, two frames (slices or sections of slices) were selected randomly and segmented manually.

These two frames are used as training input for the convolutional neural network (CNN) using the default CNN architecture: U-net with a depth of 5 layers and 64 convolutional filters per layer. After the first DL model completed the training, predictions are generated on new frames and corrections are applied on these frames when needed. These corrections add more information to the second and final CNN training for the model (22). The final model trained for an additional 100 epochs. The “early stopping” option was selected while training, that is, if the model is not improving for 10 consecutive epochs the training is aborted to review the training results. In all of the scans for both models it took two training sessions until satisfactory and accurate segmentation of the three labels was achieved.

Thresholded datasets using the mean threshold (MT) were loaded onto Dragonfly software for 3D image analysis using connectivity. Connected component refers to a set of voxels that are connected to each other. Connected component labeling marks each connected component with a distinctive label. For both segmentation methods (MT and DLM) we extracted 6-connected components and thus counted all labels with volumes between 10 and 2000 μm^3^, because objects smaller than 10 μm^3^ are at the resolution limit of the scan and are therefore subjected to significant errors. Objects larger than 2000 μm^3^ are channels and cavities not related to lacunae. See reference (23).

#### Lacunae morphometric analysis

For morphological analysis of OL we used (MT) labels with volumes in the range of 20-2000 μm^3^.

For OL concentrations we analyzed two duplicates derived from two different animals for each of the 4 bones. OL concentration was defined as the number of segmented labels in 1mm^3^. For morphometric analysis we obtained the volume and the aspect ratio of the OL labels directly from the label analysis option in Dragonfly, and the aspect ratio ψA (0 < ψA≤ 1) was determined by dividing the minimum orthogonal Feret diameter and maximum Feret diameter. The aspect ratio provides a good indication of the extent of elongation of the lacunae.

### Staining of cell nuclei

#### Bone clearing

After extraction of the petrous bones, all samples were washed with phosphate buffered saline (PBS) three times for 30 min each, then fixed in 4% paraformaldehyde (PFA) overnight. Bone clearing was carried out using the PEGASOS method (24). Briefly, bone samples were immersed in 20% EDTA (pH 7.0) at 37 °C in a shaker for 4 days for the mouse petrous bone and 16 days for the pig petrous bone until complete decalcification was achieved. To remove excess EDTA, samples were then washed with distilled water for 30 min. Then, decolorization with 25% w/v Quadrol (Sigma-Aldrich; 122262) for 2 days at 37 °C in a shaker. Samples were then placed in tert-butanol (tB) (Sigma-Aldrich; 471712) and 3% w/v Quadrol for delipidation for 2 days with gradual increasing of (tB) concentration (30%, 50% and 70% v/v). Then 70% v/v tert-butanol with 30% v/v polyethylene glycol (Sigma-Aldrich; 447943) (tB-PEG) and 3% w/v Quadrol solution was used for 2 days for dehydration. Samples were then immersed in 75% v/v benzyl-benzoate (Sigma-Aldrich; B6630) with 25% v/v polyethylene glycol (BB-PEG) medium at 37 °C for at least 1 day for clearing. All samples were preserved in freshly prepared BB-PEG clearing medium (refractive index R.I.=1.54) at room temperature in the dark.

#### Nuclei staining

samples were submerged in a DRAQ5 solution dissolved in PBS (1:1500) for two days with gentle shaking, in order to stain the osteocyte lacunae nuclei, and stained samples were then placed back into BB-PEG clearing medium (25, 26).

3D images of cleared bone were acquired using a light-sheet microscope (Ultramicroscope II, LaVision Biotec) operated by the ImspectorPro software (LaVision BioTec). The microscope is equipped with an Andor Neo sCMOS camera (2,560 × 2,160, pixel size 6.5 μm x 6.5 μm) 16 bit, and an infinity corrected setup 4X objective lens: LVBT 4X UM2-BG (LVMI-Fluor 4X/0.3 Mag. 4x; NA: 0.3; WD: 5.6-6.0 mm), with an adjustable refractive index collar set to the refractive index of BB-PEG (1.54). The light sheet was generated by scanning a supercontinuum white light laser (emission 460 nm – 800 nm, 1 mW/nm – 3 (NKT photonics). The following excitation band pass filters were used: 470/40 nm for tissue autofluorescence – providing general morphology. 617/83nm for DRAQ5. The light sheet was used at 80% width and maximum NA (0.154). Laser power of 80 %. The emission filters used were: 525/50 for autofluorescence, 690/50 for DRAQ5. Stacks were acquired using 5 μm step-size and a 200ms exposure time per step. Imaris (Bitplane) was used to create 3D reconstructions and animations of the bone.

## Results

The experimental strategy that we used was to compare two different layers of the petrous bone, one adjacent to the otic cavity (inner periosteal layer) and one close to the outer surface (outer periosteal layer). These layers were then compared to another bone in the skull complex (the temporal bone). See figure 1 for locations, and S.I. Movie.1 showing a video of the petrous and temporal bone complex. We also compared these bones to cortical bone from pig femora. Osteocyte lacunae concentrations have been studied in other animal femora (27, 28).

**Figure 1:**
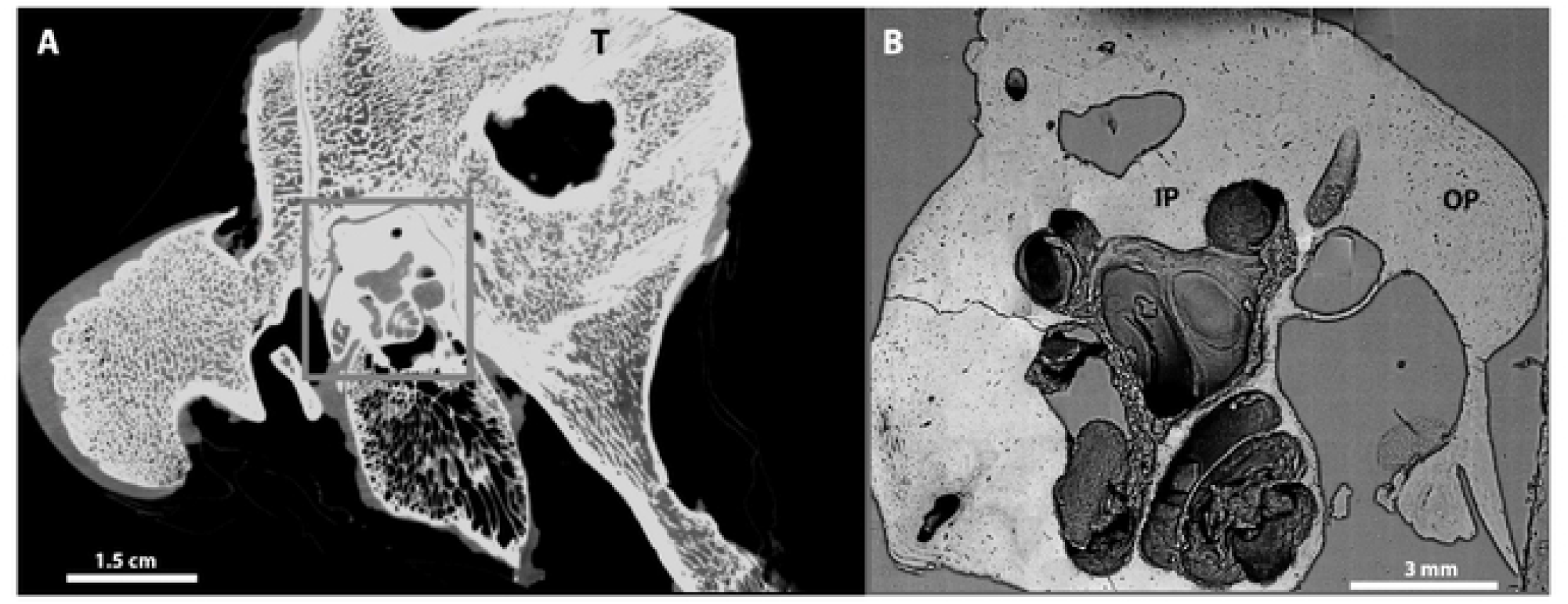
(A) A slice from a low-resolution μ-CT tomogram obtained from the pig right temporal bone before extraction of the petrous bone showing the otic chamber in the grey rectangle. (B) SEM backscattered electron image automated image map of a transverse section of a pig petrous bone after extraction, sectioning and polishing. The temporal bone samples were collected from the same location (T) from two different pigs (Figure 1A). The locations of petrous bone samples from the outer and inner periosteal layers (OP and IP) are shown in Figure 1B. These samples were collected from the same locations in two different pigs.

One high resolution micro-CT tomogram was obtained from each of the 8 samples: four from two petrous bones (two from the inner (IP) and two from the outer (OP) periosteal layers), two from temporal bones (T) and two from femora (F). The sub-micrometer voxel size of each μ-CT scan (Table 1) enabled the reliable detection of all osteocyte lacunae from all the different bone tissues (23, 29). Osteocyte lacunae (OL) were counted and measured from all sample scans following segmentation and analysis. Figure 2 shows low and high magnification views of a μ-CT slice from the inner petrous bone (IP) sample with the lacunae segmented according to the two methods (Mean Threshold (MT) and the Dragonfly deep-learning model (DLM)). Upon close examination of individual lacunae, it can be seen that the DLM segmentation labels cover a slightly larger area of OL than the MT segmentation labels (Figure 2). Table 1 shows the OL concentrations per mm^3^ of bone volume for both methods from all of the 8 samples. We used the same range of lacunae volumes that was used by (23) namely 10-2000 μm^3^, so that we could compare our results with theirs. However lacunae with volumes between 10-20 μm^3^ are very small and quite abundant. We therefore also used a range of 20-2000 μm^3^ for the MT method. The results are shown in Table 1 and plotted in Figure 3. Despite the different ranges and segmentation methods used, it is clear that both IP samples contain significantly more OL than the other bone samples. In fact the IP samples contain about 2-fold more OL than the OP samples, and about 3-fold more OL than the femora.

**Table 1:** Shows the voxel size and scan volume (Vol.) for all μ-CT scans and the concentrations of OL from all samples. The mean threshold (MT) segmentation results are represented for OL concentration in two OL volume ranges: one for OL volumes between 10-2000 μm^3^ (the range used by Carter *et al*. (23, 30)); and 20-2000 μm^3^. Table 1 shows one range for the deep-learning model (DLM) segmentation method for OL volumes between 10-2000 μm^3^

| Sample | Voxel size<br>μm<sup>3</sup> | Scan Vol.<br>mm<sup>3</sup> | MT (10-2000μm<sup>3</sup>)/mm<sup>3</sup> | MT (20-2000μm<sup>3</sup>)/mm<sup>3</sup> | DLM (10-2000μm<sup>3</sup>)/mm<sup>3</sup> |
| --- | --- | --- | --- | --- | --- |
| IP2 | 0.9 | 0.5 | 106000 | 97000 | 85000 |
| IP3 | 0.9 | 0.6 | 110000 | 99000 | 94000 |
| OP2 | 0.9 | 0.5 | 59000 | 55000 | 43000 |
| OP3 | 0.8 | 0.4 | 77000 | 60000 | 52000 |
| T2 | 0.8 | 0.4 | 85000 | 46000 | 25000 |
| T3 | 0.9 | 0.5 | 58000 | 45000 | 30000 |
| F1 | 0.8 | 0.3 | 38000 | 28000 | 27000 |
| **F2** | 0.8 | 0.3 | 34000 | 30000 | 29000 |

**Figure 2:**
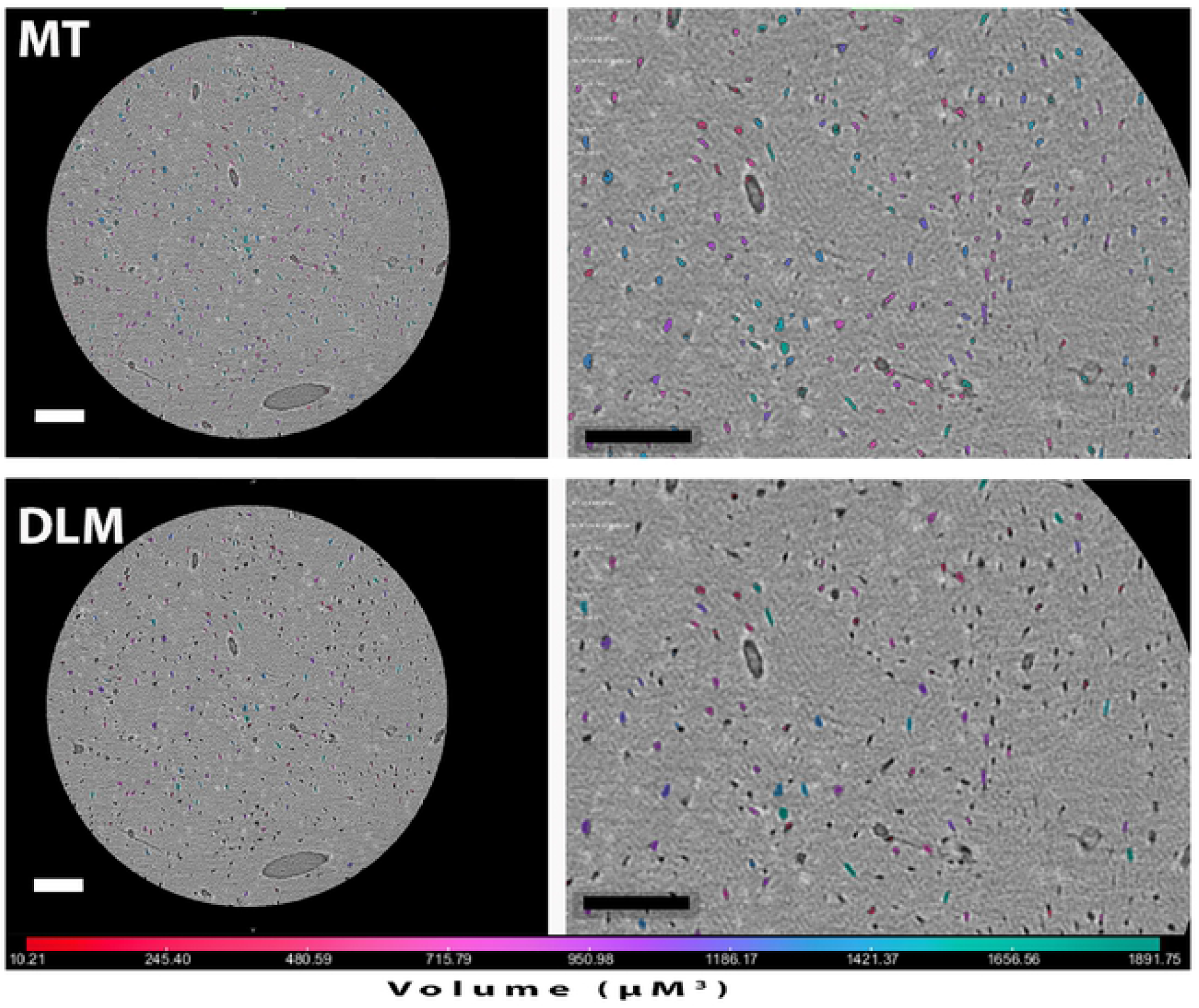
High resolution μ-CT tomogram slice of the inner periosteal layer sample (IP2) after MT (top) and DLM (bottom) segmentation. Left images show the whole field of view at the same slice location, and right images show insets of the OL labels as obtained from the two different segmentation approaches. Colors are a qualitative indication of the volume of the OL labels. Scale bars =100μm.

**Figure 3:**
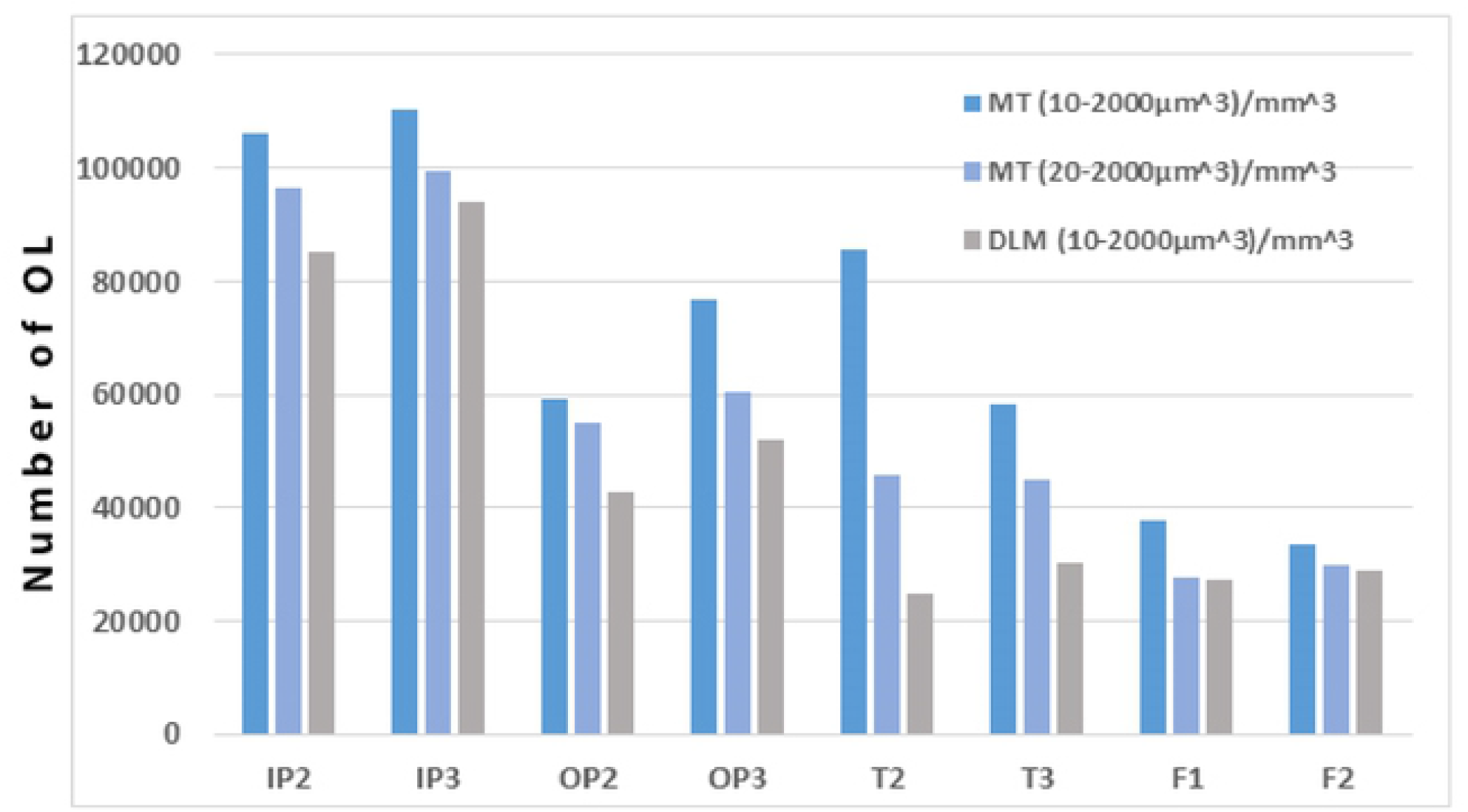
The OL concentration using the two segmentation approaches. Dark blue represents the MT segmentation with volumes from 10-2000μm^3^, light blue is the same segmentation method but with volumes from 20-2000 μm^3^, and grey represents the DLM segmentation for OL volumes between 10-2000μm^3^.

The next question we addressed was whether or not the OL in the 4 different bone samples have different volumes and/or shapes. Table 2 shows the average OL volumes and average aspect ratios obtained from the MT segmentation for labels between 10 and 2000 μm^3^. As the volume standard deviations are very large, we also plotted the distributions of OL as a function of volume (Figure 4). The OL in the inner periosteal layer (IP) are smaller than those in the outer periosteal layer (OP). The IP lacunae have similar volumes to those in the femora. The large standard deviation and the unusual distribution of T2 (table 2), preclude concluding anything about its volume distribution. The aspect ratio values show no significant difference between all 8 samples.

**Table 2:** OL average volume (Avg. Vol.) and average aspect ratio (Avg. AR) measurements of osteocyte lacunae segmented labels, together with their standard deviations (Stdev.). The data shown are obtained from the mean thresholding segmentation method (MT) for OL volume ranges between 20 and 2000 μm^3^.

| Sample | Avg Vol. ($\mu\text{m}^3$) | Vol. Stdev. ($\mu\text{m}^3$) | Avg. AR | AR Stdev. |
| --- | --- | --- | --- | --- |
| IP2 | 128 | 81 | 0.41 | 0.11 |
| IP3 | 140 | 117 | 0.51 | 0.11 |
| OP2 | 404 | 278 | 0.47 | 0.11 |
| OP3 | 162 | 105 | 0.45 | 0.12 |
| T2 | 166 | 190 | 0.49 | 0.11 |
| T3 | 212 | 155 | 0.45 | 0.12 |
| F1 | 122 | 72 | 0.34 | 0.10 |
| F2 | 137 | 112 | 0.39 | 0.11 |

**Figure 4:**
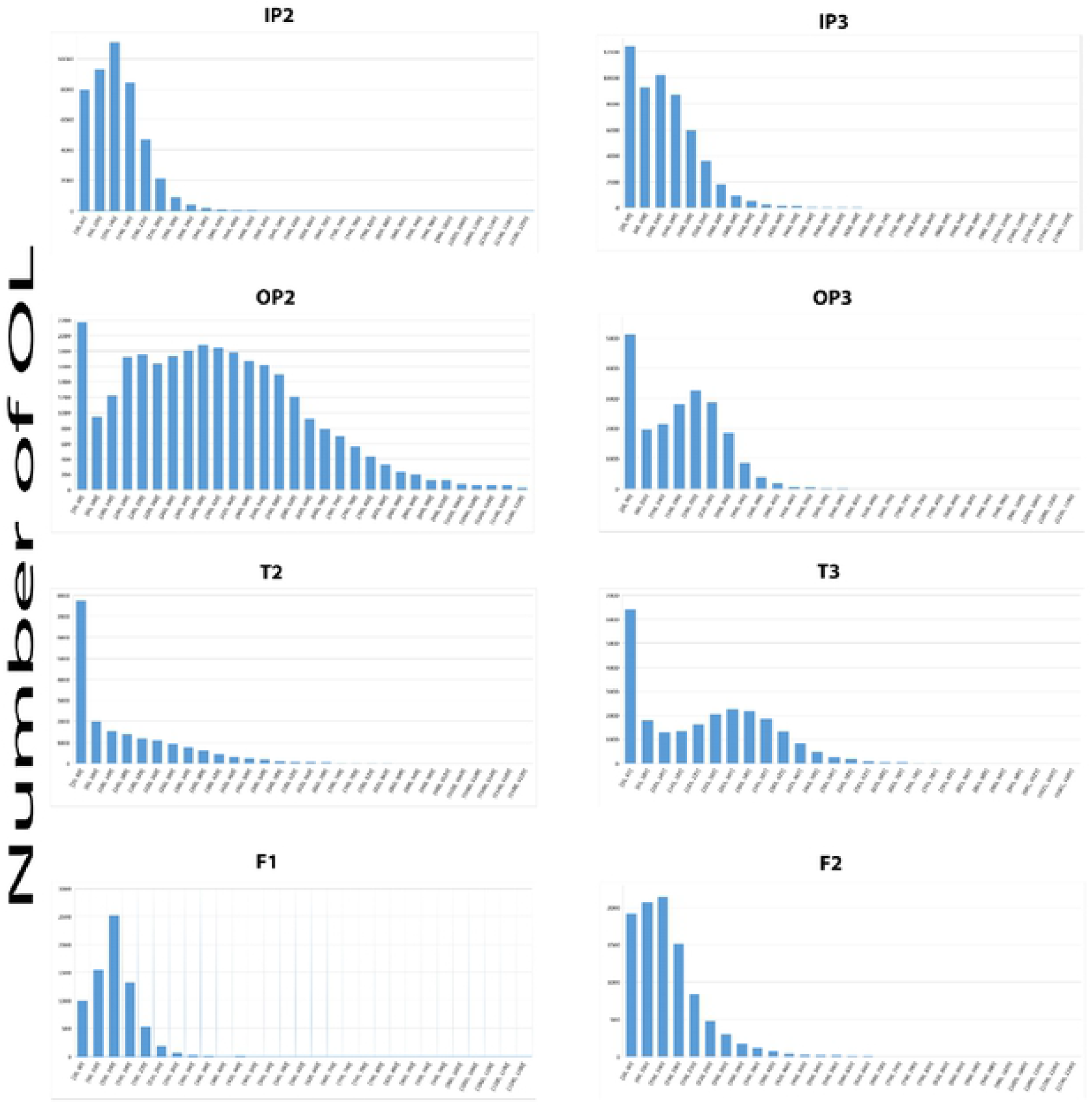
OL volume histograms from all the samples showing the OL volume distribution from 20 to 1200 μm^3^. OL volume average and standard deviations are provided in Table 2. Note that the unusual distribution of T2 may be due to the poor resolution of the scan (see Methods).

To check the viability of osteocytes in a section containing the outer and inner periosteal layers, we used DRAQ5 to stain the cell nuclei of a cleared and fluorescently dyed petrous bone from pig. As controls we also studied mouse bones as the method was developed using mouse bones (see S.I. Figure 1). The autofluorescence signal from the sample is thought to be derived mainly from collagen (green). Figure 5 shows light-sheet images obtained in two channels. The DRAQ5 nuclear staining shows the presence of nucleated osteocytes in the outer-inner periosteal bone layers outside the cochlea (red). The lacunae in the outer layers of the petrous bone therefore do contain osteocytes together with the DNA in their nuclei.

**Figure 5:**
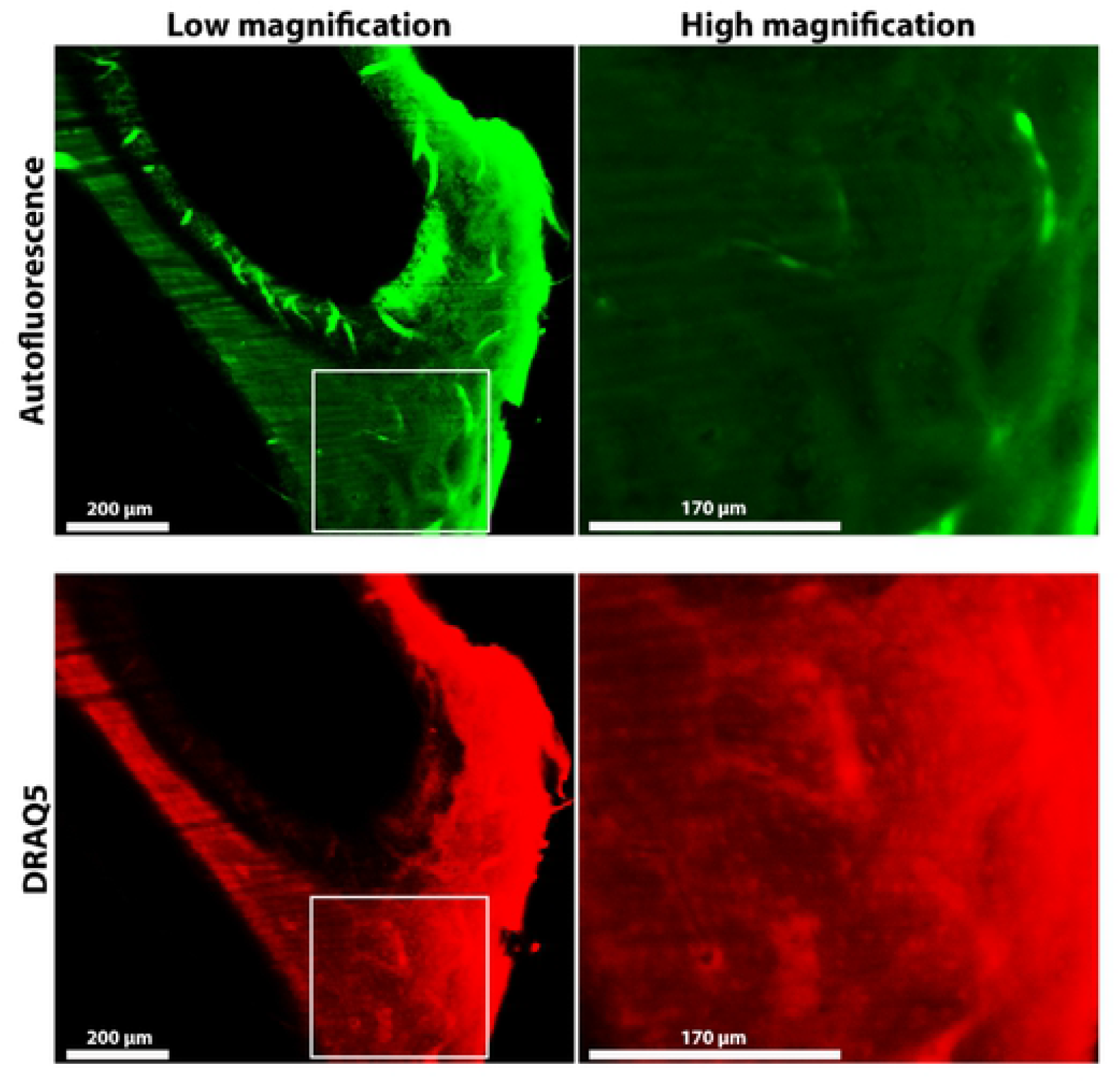
Light-sheet fluorescent microscopy images of a cleared pig petrous bone. All images are from the same Z-depth with (green) autofluorescence and (red) DRAQ5 signals. Low magnification images show the cleared bone tissue from the cochlear cavities towards the outer bone surface. High magnification images of areas indicated by the rectangles are shown. The local concentrations of DRAQ5 sained nuclei can best be seen in the high magnification image.

## Discussion

This study shows that the inner periosteal layer (IP) has significantly more osteocyte lacunae than the outer periosteal layer, and both these layers have more osteocyte lacunae than the femora. The temporal bone OL concentrations are less reliable but are certainly well below the concentration in the IP. Furthermore, we show that the petrous bone does contain concentrations of DNA, which are from osteocytes within the lacunae.

We can compare the OL concentrations and volumes that we obtained from the pig femora to concentrations reported in the long bones of other species. Carter *et al*. measured the OL concentrations in a 20 year old human male femur. The OL concentrations ranged between 26,000 and 37,000 per mm^3^ of bone (23). Androwski *et al*. obtained 22,000 per mm^3^ from human cortical bone (20). We obtained similar values from cortical pig femora. The lacunar volumes measured in the human male are around 400 μm^3^, whereas in the human female group they are on average around 240 μm^3^ (23, 30). Our measurements of the two pig femora OL volumes are on average 120 and 140 μm^3^ and are therefore considerably smaller than the human bone values reported by Carter *et al*. (23, 30).

These observations show that modern pig petrous bone, and in particular the part of the petrous bone near the otic chamber (IP layer), contains at least three times as much DNA as the femoral bones of the pig. Thus as the petrous bone initially has much more DNA, during fossilization (diagenesis) the petrous bone may also preserve more DNA. This therefore could be one reason why fossil petrous bones contain more aDNA than other bones (8, 9, 11), assuming petrous bones of other animals, including humans, also contain more osteocytes. It would therefore be interesting to carry out a systematic study of osteocyte lacunae concentrations in bones of the human skeleton, as other bones may also have high concentrations of lacunae and hence possibly high concentrations of aDNA in the same fossil bone. It would also be of interest to carry out a detailed 3D study of the structure of the petrous bone, as this may reveal other structural properties that might be responsible for the good preservation of aDNA. We are currently carrying out such a study using focused ion beam – scanning electron microscopy (FIB SEM).

It has been proposed that the higher density of the petrous bone is responsible for the good preservation of aDNA (8). It is known that DNA has a high affinity for carbonated apatite and that bound DNA is less susceptible to enzyme degradation (31, 32). However higher bone density results in a much lower surface area (33) and hence probably less DNA binding. It has also been suggested that the high density of petrous bone reduces bacterial activity and therefore post-mortem decay of DNA (8). It is not clear why higher density per se should reduce bacterial activity. A reduction in bacterial activity may be the consequence of some unique structural features of the petrous bone, such as the characteristics of blood vessels and canaliculi.

It is interesting to consider the possible functions of having so many osteocytes in the petrous bone close to the otic chamber. Doden and Halves postulate that the otic capsule bone has a physiological function related to sensation (17). Aarden *et al*. summarize the functions of osteocytes as follows: 1) communication bridges between inside and outside the bone tissue; 2) increasing the mineral surface exposed to extracellular fluid and cellular activity; 3) providing a repair mechanism in addition to remodeling (34). While others show that osteocytes remodel bone matrix through perilacunar/canalicular remodeling (PLR) and they highlight the importance of osteocytes for improving bone fragility (35, 36). The ability of osteocytes for mechanosensing is well documented (37). Interestingly, Fung *et al*. exposed osteocytes to low intensity pulsed ultrasound (LIPUS). This stimulated the osteocytes and promoted the mechano-transduction between osteocytes and osteoblasts (38). Therefore, it is plausible that the high concentration of osteocytes in the petrous bone, especially in the layers closer to the otic capsule, might be a result of bone adaptation for improved acoustical/sensing.

## Conclusions

We show that the petrous bone of the pig has three-fold more osteocyte lacunae, than the femoral compact bone and hence the petrous bone has significantly more DNA occluded inside the bone. This might be one factor that leads to the preservation of high amounts of DNA in fossil bones. We raise the possibility that osteocytes of the petrous bone might have acoustical functions.

## Acknowledgements

We thank the Kimmel Center for Archaeological Science for partial funding of JI’s fellowship, as well as the analytical costs. We also thank Alejandro Aguilera Castrejon from the department of Molecular Genetics in Weizmann Institute of Science for his help in this project.

## Supporting Information Captions

S.I. Figure 1: Fluorescent microscope images at the same Z-depth of mice petrous bone after total clearing using the PEGASOS protocol (24) and DNA staining using DRAQ5. (A) Autofluorescence signal (green), (B) DRAQ5 signal (magenta), and (C) both signal channels combined. In (B and C) the DRAQ5 staining of the spiral ganglion (S.G.) and the inner membranes of the cochlea (Coc) clearly show high concentrations of DNA in the mouse petrous bone. This confirms that the protocol used for the petrous bone should be reliable, even though the petrous bone takes much longer to clear. Scale bar = 500μm.

S.I. Movie 1: Low resolution micro-CT scan of the extracted skull bone from pig showing the petrous, temporal, tympanic bullae, and zygomatic arch bones. Note that the petrous bone contains no spongey bone component. Pixel size = 83μm and scale bar = 1cm.

